# Double Machine Learning with Multi-Gene Shared Background for Causal Inference in Single-Cell Data: Grouping Deviation Follows a Random Walk and the Accuracy-Compute Trade-Off

**DOI:** 10.64898/2026.08.08.743701

**Authors:** Wei Ye, Xinyu Jiang, Feng(Ben) Shen

## Abstract

In high-throughput single-cell transcriptomics (*p* ≈ 20,000 genes), performing double machine learning (DML) causal inference on *q* ≈ 5,000 target genes requires nuisance function fits that grow linearly with the number of targets (*K_f_* cross-fitting folds, *K_f_* = 5 or 10), far exceeding feasible computational budgets, especially with deep learning. We propose a Randomized Partition Strategy (RPS): randomly divide target genes into groups, share one background compression per group, reducing deep learning model training to *q*/*m* runs (*m* = group size) --- a factor of *m* savings. The cost of grouping is accuracy loss --- we prove that the cumulative deviation of the estimator follows a one-dimensional drift-free symmetric random walk, with diffusion variance growing linearly with group size and mean squared displacement equaling the mean squared error, so accuracy loss is predictable: *m* = 1 is always optimal, accuracy cost is monotonically increasing, and a small accuracy sacrifice yields *m*-fold compute savings. On GSE189050 SLE single-cell data (Memory B cells, *n* = 2120), both PCA and DL methods converge to the same conclusion, confirming the random walk mechanism is method-independent; an unexpected finding is that DL diffusion growth is only 16%, far slower than PCAs 7.4 times. This work provides a quantifiable theoretical foundation for compute strategy selection in single-cell high-dimensional causal inference.

## 1. Introduction

### 1.1 Background

Causal inference is a central challenge in modern data science. Double machine learning (DML), as a framework for causal inference with high-dimensional nuisance variables, traces its roots to semiparametric estimation of partially linear models ^[1]^: it uses machine learning to estimate nuisance functions, constructs an orthogonal score via Neyman orthogonality that is insensitive to nuisance estimation errors, and employs cross-fitting to prevent overfitting of nuisance functions from contaminating the target parameter estimate, yielding √n-consistent and asymptotically normal estimates of the target parameter ^[2]^. This method has been widely applied in economics, epidemiology, and social science. In recent years, with the development of high-throughput biotechnology, causal inference methods have begun to be applied to large-scale data at the molecular biology level. Applying causal machine learning to single-cell transcriptomics has become an active direction ^[3][4]^: de-confounding double machine learning variants have been used for robust causal estimation under high-dimensional gene expression ^[5]^, and a robust causal framework combining semiparametric machine learning under unmeasured confounding (causarray)^[6]^ has also been proposed; causal identification methods for single-cell perturbations (e.g., CINEMA-OT)^[7]^ and perturbation effect estimation methods based on causality-aware generative models and variational causal inference (scCausalVI)^[8][9]^ have emerged, the assumptions and methods for causal effect estimation in single-cell CRISPR screens have also been systematically reviewed ^[10]^, and trajectory-inference-based single-cell Mendelian randomization extends causal gene identification to dynamic phenotypic differences (ti-scMR)^[11]^; unified perturbation data benchmarks (scPerturb)^[12]^ provide standard datasets, highlighting the practical need for efficient causal estimation in ultra-high-dimensional, large-target-set settings. Among these, a DML pipeline employing shared unsupervised deep learning to construct background nuisance functions has been applied to causal effect analysis in single-cell transcriptomics data ^[13]^, providing a feasible path for de-confounding inference in high-dimensional gene expression settings.

Consider the typical single-cell scenario: a researcher has dozens of target genes (e.g., all genes in a signaling pathway) or even thousands (e.g., all genes in the transcriptome), and needs to estimate the causal effect of each target gene on a binary outcome (such as cell type differentiation state or drug response) while controlling for other gene expression levels.

The key characteristic of this scenario is simple: each target gene requires its own independent model estimation --- *q* target genes means *q* complete DML pipelines, with computational cost growing linearly with the number of targets.

Before formalizing the problem, we first outline the core intuition of DML in plain language, serving as an intuitive anchor for the discussion that follows. Consider the fundamental question: is *T* truly a cause of *Y*? The central challenge of DML is the presence of another variable *X* that simultaneously affects both *T* and *Y* (confounding), so the arrow *T* → *Y* may be polluted by spurious associations created by *X*. DML’s strategy is to “remove the influence of *X* on both sides” --- construct residuals *T*′ and *Y*′ that cannot be explained by *X*, and then read off the clean causal effect from the relationship between *T*′ and *Y*′. This idea underpins all derivations and experiments in this paper; the next section provides a formal version.

The key to the DML pipeline lies in how the nuisance function (*E*[*T*|*X*], *E*[*Y*|*X*]) is constructed. Fig. 1 uses three panels to summarize the compute logic of three construction strategies. Panel A is the standard DML approach: each target gene is fitted independently with a complete fit, with model fits ≈ number of target genes *q* times the number of cross-fitting folds *K_f_*. Panel B is the approach of a recent preprint ^[13]^: equivalent to standard DML, but the two nuisance functions share one unsupervised deep compression representation, with model fits ≈ *q*, eliminating the *K_f_* cross-fitting folds. Panel C is the approach proposed in this paper: further reducing model fits by sharing one model fit across multiple target genes --- multiple targets share one model fit. However, Panel C inevitably introduces bias; precisely how much bias is introduced is the main question this paper investigates. The following explains why bias arises.

**Figure 1:**
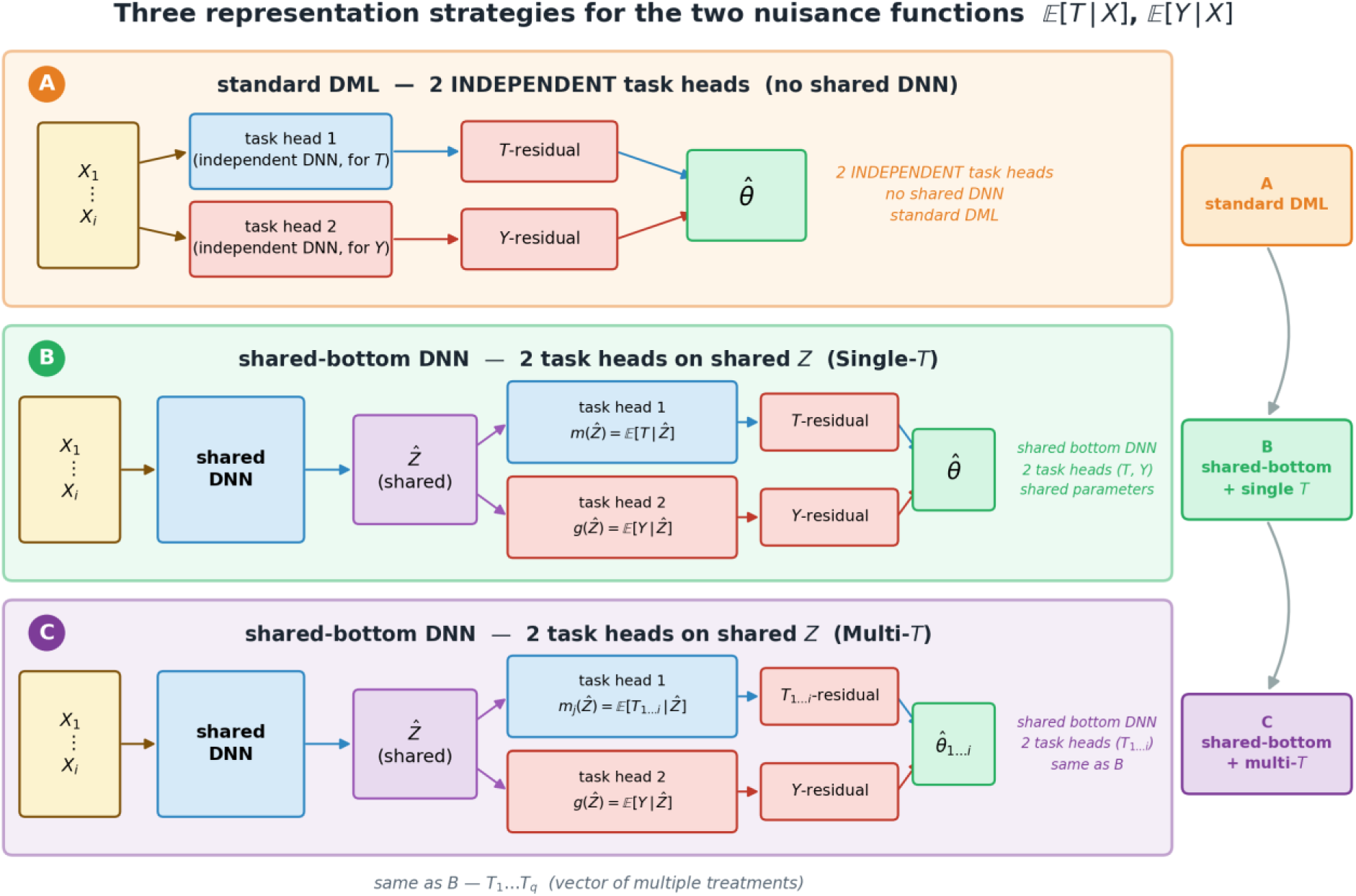
Three strategies for constructing nuisance functions in DML (conceptual illustration). Panel A (dual independent heads, standard DML): each target uses two independent networks to estimate *E*[*T*|*X*] and *E*[*Y*|*X*] separately, with no DNN sharing --- model fits ≈ number of target genes *q* times the number of cross-fitting folds *K_f_*. Panel B (shared compression, preprint method): a shared DNN compresses high-dimensional *X* into low-dimensional *Z*, with two linear heads *m*(*Z*) = *E*[*T*|*Z*] and *g*(*Z*) = *E*[*Y*|*Z*] on top of *Z* --- equivalent to standard DML, with model fits ≈ *q*, eliminating the *K_f_* cross-fitting folds. Panel C (shared compression, multi-target, this paper): a shared *Z* with *q* sets of task heads *m_j_*(*Z*) = *E*[*T_j_*|*Z*] and *g*(*Z*) = *E*[*Y*|*Z*] --- multiple targets share one deep compression, reducing model fits to *K* = *q*/*m*; however, group-mates become omitted confounders for each other, inevitably introducing bias (see Fig. 3 for the bias mechanism).

#### We adopt standard DML notation

*T* (treatment variable, in this paper the j-th target gene *T_j_*), *X* (all control/confounding variables), *Y* (outcome); *X_j_* denotes the subset of *X* excluding *T_j_*.

Why does grouping cause bias? Fig. 3 directly illustrates the mechanism from a variable-set perspective: when *m* = 1, target *T*_1_ is in its own group with all other target genes entering the background as controls --- no omitted confounders, unbiased estimation, corresponding to the random walk starting point *S*(1) = 0; when *m* = 3, *T*_1_, *T*_2_, *T*_3_ are grouped together and excluded, becoming omitted confounders for each other --- the estimate for *T*_1_ is randomly shifted by the effects of *T*_2_ and *T*_3_. Each additional group-mate adds one step; *m* = 3 is equivalent to *m* − 1 = 2 steps. This causal chain --- group-mates becoming omitted confounders → random shift in estimates → accumulation into a random walk --- is precisely the statistical mechanism that Section 2.1 will systematically characterize.

Since grouping inevitably introduces bias, the core question this paper addresses is: How large is this bias? What law does it follow? Can we predict it before grouping? Fig. 4 provides a visual summary of this trade-off: the upper half (compute dimension) shows that *m* = 1 requires *q* autoencoder training runs, *m* = 3 only *q*/*m* --- saving a factor of *m*; the lower half (accuracy dimension) shows that at *m* = 1, all target gene estimates are tightly centered on the true value *θ* (*S*(1) = 0, MSD = 0); at *m* = 3, estimates spread around the true value according to *Var*[*S*] ∝ (*m* − 1) --- a small accuracy loss is the price paid for computational savings. Trading a small accuracy loss for order-of-magnitude compute savings is the central proposition of the RPS framework.

From Fig. 2-4, we see that shared background saves computation but inevitably introduces bias. The magnitude and law of this bias are not immediately obvious --- they depend on a complex interaction of factors including the correlation, effect size, and sample size of the excluded group-mates. Section 2 will systematically characterize this bias --- we show that the cumulative deviation 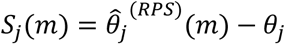 of the grouped estimator follows a one-dimensional random walk, with diffusion (variance) proportional to *m* − 1 and generally drift-free, providing a quantifiable mechanistic explanation for the accuracy cost. This random walk characterization not only answers the question “how large is the bias?” but also places the decision criterion of “choose *m* based on the accuracy/compute budget” (Section 2.2.3) on a rigorous statistical foundation rather than empirical rule-of-thumb.

**Figure 2:**
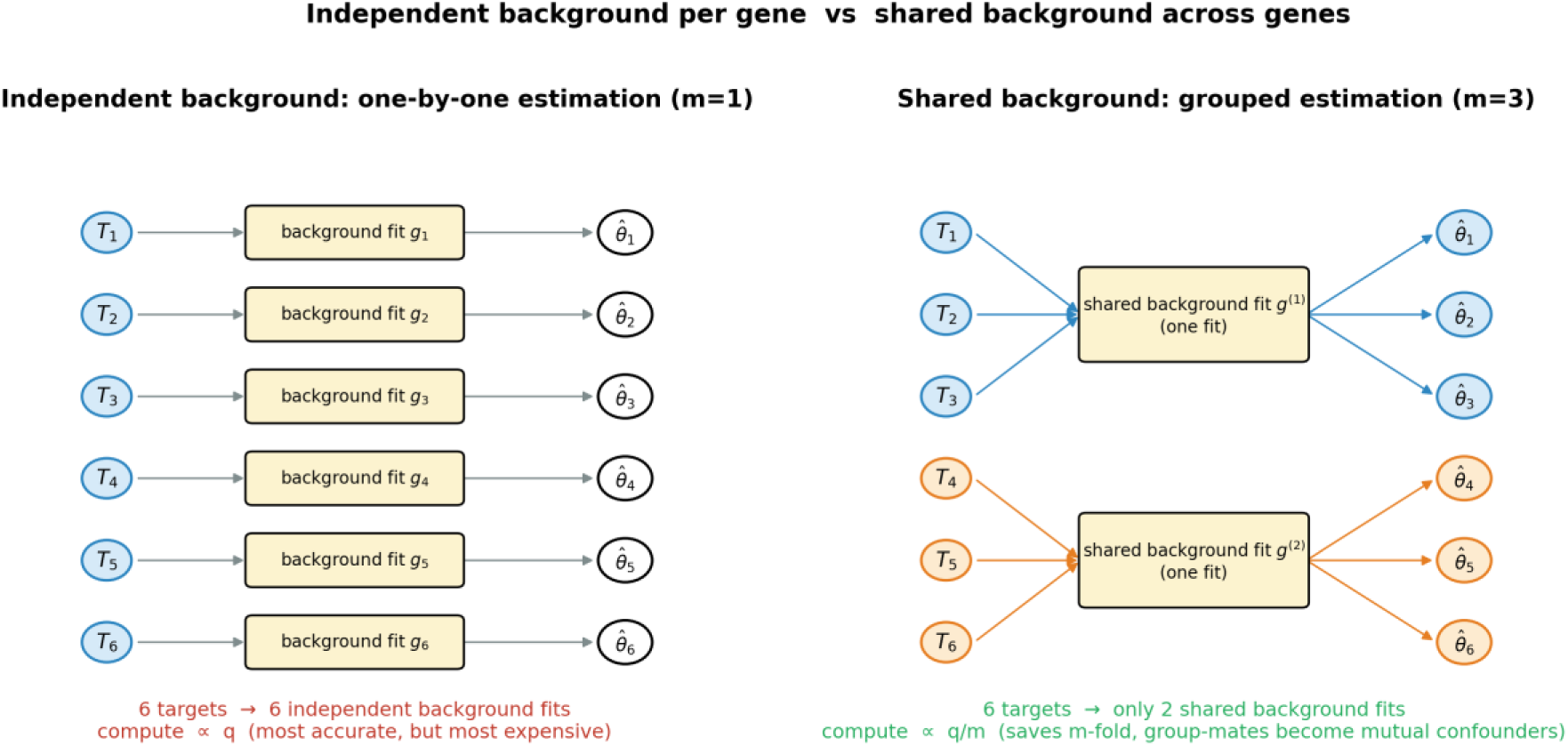
Comparison of individual estimation (*m* = 1) and grouped estimation (*m* = 3) (conceptual illustration). Left: independent background, individual estimation (*m* = 1) --- each target gene *T_j_* fits its own background *g_j_* independently, for a total of *q* independent redundant fits; statistically optimal (corresponding to the gold-standard starting point *S*(1) = 0 of the random walk), but with computational cost ∝ *q*. Right: shared background, grouped estimation (*m* = 3) --- every *m* target genes share one background fit *g*^(*k*)^, requiring only *q*/*m* fits; saving a factor of *m* in computation, but group-mates are excluded and become omitted confounders for each other (see Fig. 3), causing bias in the estimates. RPS is the systematic version of the latter approach.

**Figure 3:**
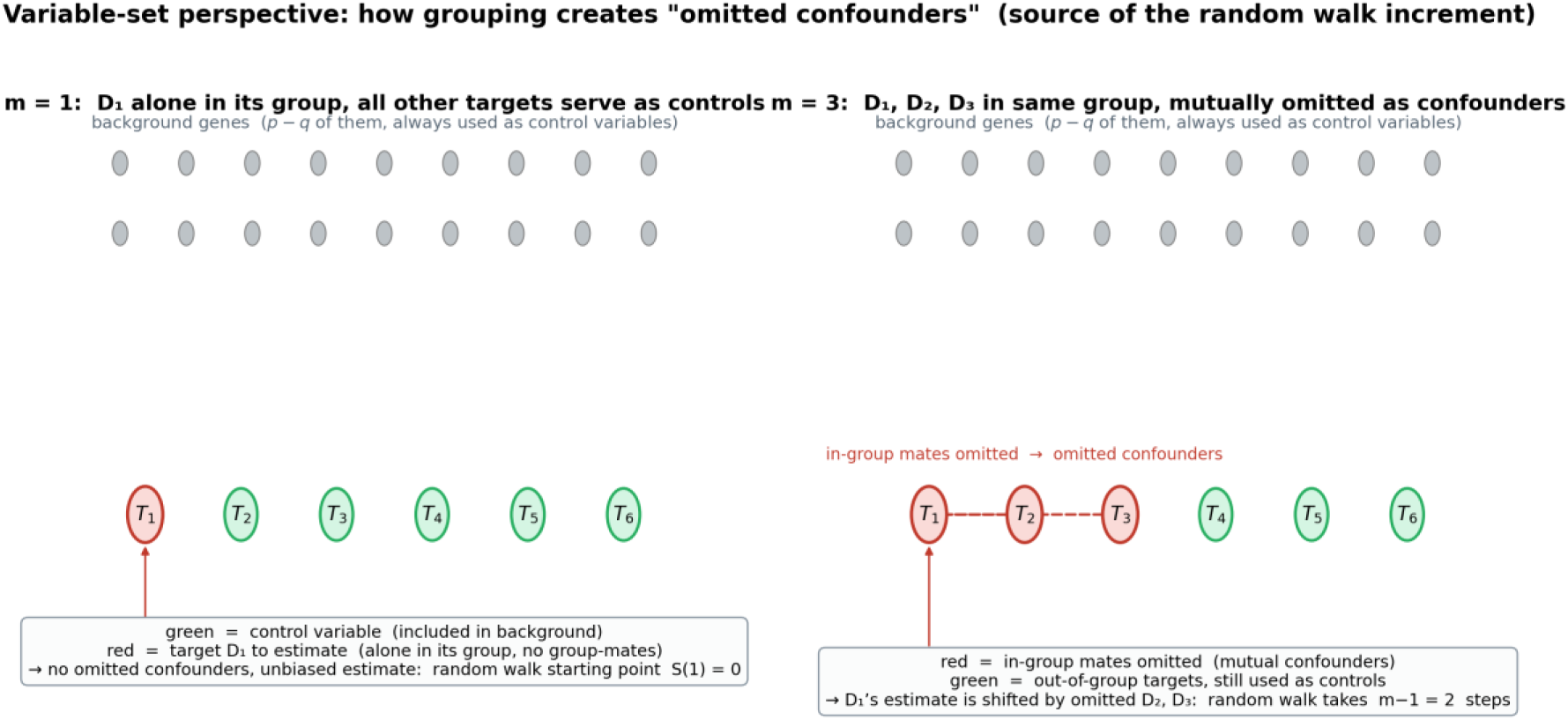
How grouping creates omitted confounders --- the source of random walk increments (variable-set perspective, conceptual illustration). Left: *m* = 1, *T*_1_ alone in its group, all other targets serve as controls. Target *T*_1_ (red) is alone in its group; *T*_2_, …, *T*_6_ (green) together with all background genes (gray grid above) enter the control variable set --- no omitted confounders, unbiased estimation, corresponding to the random walk starting point *S*(1) = 0. Right: *m* = 3, *T*_1_, *T*_2_, *T*_3_ grouped together and excluded, becoming omitted confounders for each other. Red *T*_1_, *T*_2_, *T*_3_ are simultaneously excluded from the control set; green *T*_4_, *T*_5_, *T*_6_ remain as controls --- *T*_2_ and *T*_3_ become omitted confounders for *T*_1_, injecting a random increment into the estimate for *T*_1_. Each additional group-mate adds one step; thus *m* = 3 is equivalent to *m* − 1 = 2 steps.

**Figure 4:**
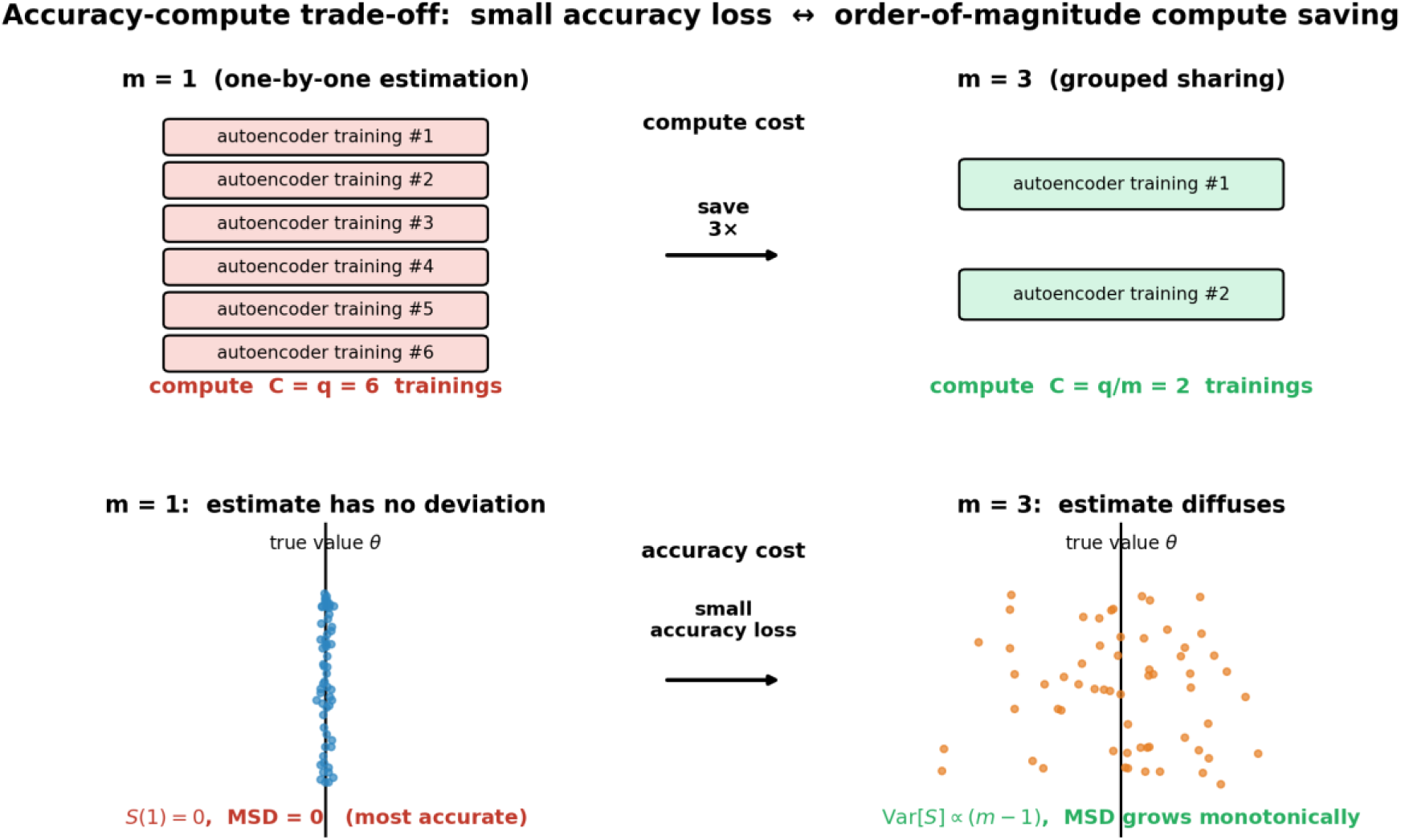
Accuracy-compute trade-off side by side (conceptual illustration). Top: compute cost --- *m* = 1 individual estimation requires training one autoencoder per target (total *q*, most expensive), *m* = 3 grouped sharing requires only *q*/*m* runs; in the deep learning scenario of this paper, this count is directly observable by the reader. Bottom: accuracy cost --- at *m* = 1, all target gene estimates are tightly centered on the true value *θ* (*S*(1) = 0, MSD = 0, most accurate); at *m* = 3, estimates spread around the true value according to *Var*[*S*] ∝ (*m* − 1) (mean squared displacement MSD increases monotonically). A small accuracy loss is the price paid for computational savings. The two rows together present the core RPS proposition: compute cost drops substantially with *m*, while accuracy cost rises slowly --- an asymmetric trade-off.

## 2. Methodological Framework

### 2.1 Random Walk Characterization of Grouping Deviation

This section unifies the bias and variance of the RPS estimator into a single dynamical picture: each time a target gene shares the same background control with the target of interest (while itself remaining uncontrolled), it becomes an omitted confounder, causing the estimate to shift randomly by one step; the shifts accumulate along the number of shared targets *m* − 1, forming a one-dimensional random walk. Since the shared companions are randomly selected and the correlations and effects have mixed signs, the expected increment per step is generally zero, hence the general case is a diffusion-dominated (drift-free) random walk. A significant drift appears only when the product of correlation and effect has a systematic same sign (e.g., co-expression modules with uniform sign). Random grouping is merely a computationally efficient implementation of “sequentially adding random shared targets“: *m* targets sharing one background estimate ⇔ each target takes *m* − 1 steps at once, reducing the computational cost from *q* to *q*/*m* (Section 2.2.2) --- the random walk is the essence of the phenomenon; random grouping is merely the means of implementation. The RPS bias follows from the standard omitted variable formula (appendix). Fix a target *T_j_*; its cumulative deviation 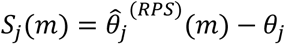 along group size *m* is a one-dimensional random walk: each additional group-mate injects a random increment 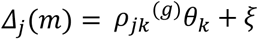. Increments are approximately uncorrelated, and variance accumulates to 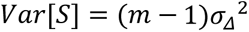 universally; the increment mean *μ_j_* = *E*[*ρ_jk_ θ_k_*] is ≈ 0 when correlations and effects have mixed signs (drift-free), and significant only when systematically same-signed. Mean squared displacement 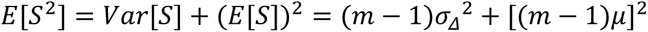 equals *MSE*(*m*), so accuracy loss is predictable.

**Figure 5:**
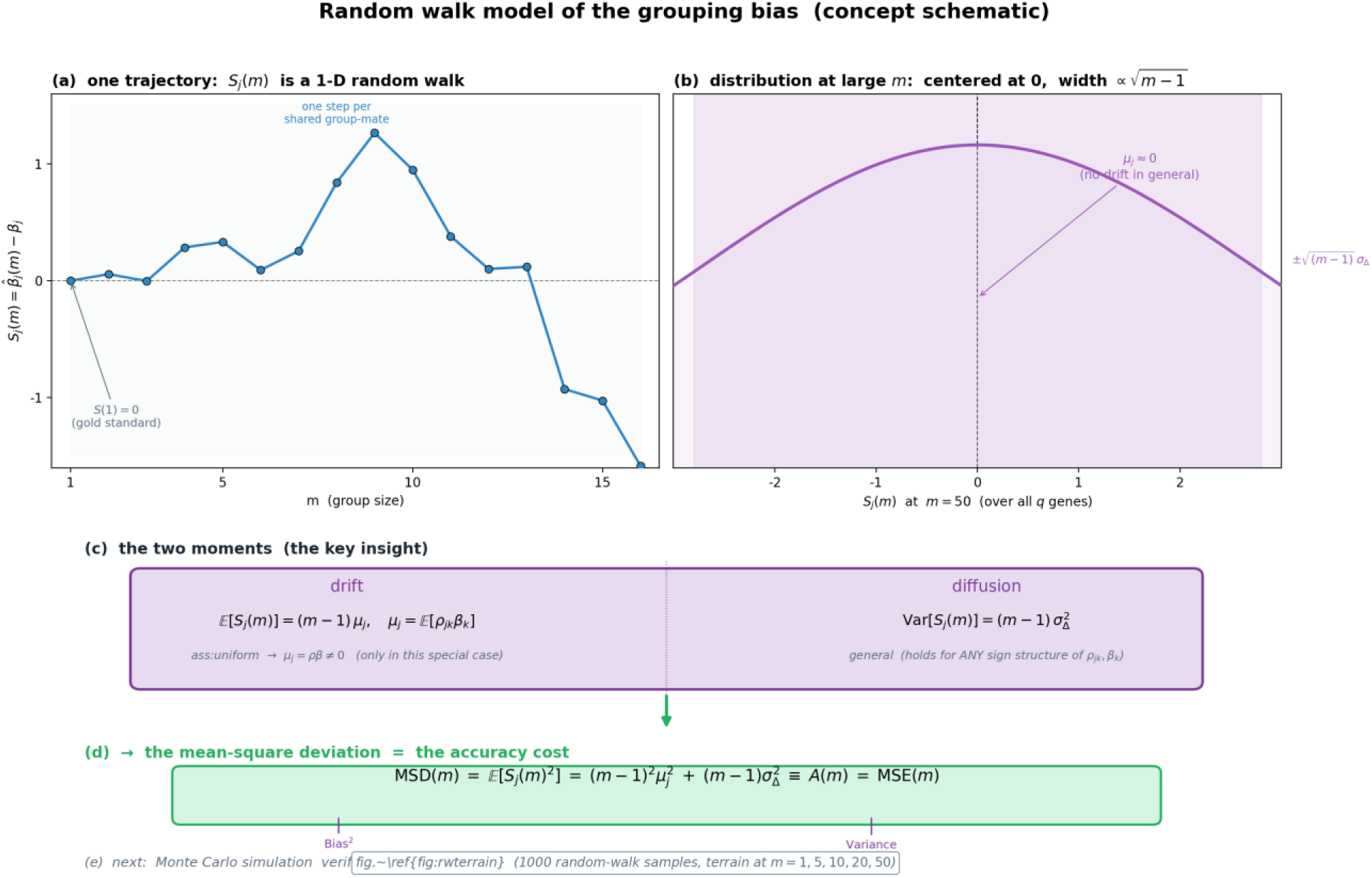
Random walk model of grouping deviation --- theoretical conceptual illustration. (a) Single trajectory: for a single target gene *T_j_*, the cumulative deviation *S_j_*(*m*) along *m* evolves as a one-dimensional random walk, with *S*(1) = 0 as the gold-standard starting point. (b) Overall distribution: in the cross-section across *q* target genes, *S* at large *m* approximates a symmetric distribution centered at 0 with width ∝ √*m* − 1 --- in the general case (mixed-sign correlation/effect), the drift term *μ_j_* ≈ 0. (c) Two moments: *E*[*S_j_*(*m*)] = (*m* − 1)*μ_j_* (only under the homogeneity assumption with *μ_j_* = *ρθ* ≠ 0), *Var*[*S_j_*(*m*)] = (*m* − 1)*σ*^2^ (universal). (d) Accuracy cost: *MSD*(*m*) = (*m* − 1)^2^*μ*^2^ + (*m* − 1)*σ*^2^ ≡ *A*(*m*) = *MSE*(*m*) --- the random walk interpretation of “bias squared plus variance.” (e) The Monte Carlo simulation in Fig. 6 (landing terrain of 1000 independent walks) provides empirical verification of this two-moment picture.

**Figure 6:**
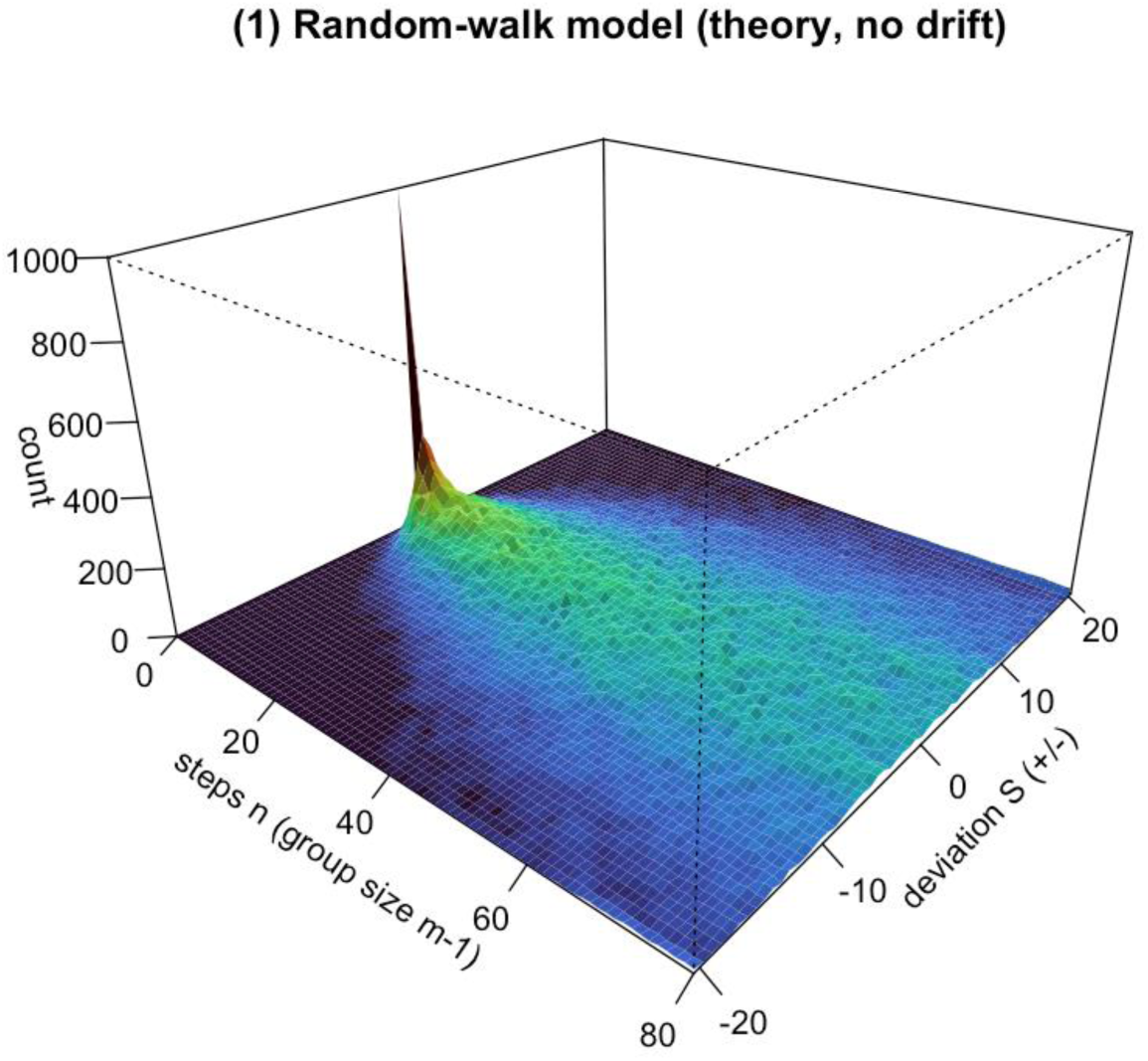
Landing terrain of a general (drift-free) random walk --- Monte Carlo visualization of the theoretical random walk model (serving as the premise to be verified by the subsequent two-layer empirical validation). We simulate 1000 independent one-dimensional random walks (each step has zero mean increment), and count the frequency of landing positions at each step: the horizontal axis is the step count *n* (= the number of other target genes sharing background control with the target of interest = *m* − 1; *m* = 1 is the 0-step gold-standard starting point, and each additional shared random target gene adds one more step), the vertical axis is the cumulative deviation *S* from the origin (± both directions), and the height (color, sqrt scale) is the landing frequency. The terrain is symmetric about *S* = 0 --- confirming that in the general case the increment mean *μ_j_* ≈ 0 with no drift; the distribution width broadens as ∝ √*n*, visually presenting the diffusion-dominated random walk picture.

### 2.2 Accuracy-Compute Trade-off and Group Size Selection

#### 2.2.1 Accuracy Cost: Monotonic Convexity of MSE

Under the linear DML setting, when the sample size *n* is sufficiently large (number of controls much smaller than *n*), the variance term changes slowly with *m*, while the bias squared term is monotonically increasing and convex. Therefore *MSE*(*m*) = *Bias*^2^(*m*) + *Var*(*m*) is monotonically increasing in *m*, with no interior minimum: *m*^∗^*_accuracy_* = *arg min*_1≤ *m*≤ *x*_ *MSE*(*m*) = 1. That is, from a purely statistical accuracy standpoint, the most precise approach is always to estimate each target variable individually. Increasing group size only monotonically sacrifices accuracy; the reason grouping is still worthwhile is that it simultaneously monotonically saves computation --- the next subsection explains why this trade-off is favorable.

#### 2.2.2 Compute Cost

In the opposite direction to accuracy cost, compute cost decreases monotonically as group size *m* increases. The reasoning is straightforward: RPS divides the *q* target variables into *K* = ⌈ *q*/*m*⌉ groups, each sharing the same control set with the most expensive nuisance fit performed once per group, requiring *q*/*m* independent estimations --- the larger the group, the fewer groups, the more compute saved. Accordingly, the compute cost (measured by the number of independent estimations) is *C*(*m*) ∝ *q*/*m* ∝ 1/*m*, i.e., the speedup factor relative to individual estimation (*m* = 1) is exactly *m*: each time the group size increases by a factor, computation decreases by the same factor. Particularly, in the single-cell + deep learning scenario of this paper, *C*(*m*) = *q*/*m* is exact: the most expensive autoencoder training is retrained for each group, with the number of training runs equal to the number of groups *q*/*m* (*m* = 1 requires *q* runs, *m* = *q* requires only 1), making *C*(*m*) a directly countable physical quantity.

#### 2.2.3 Trade-off: Small Accuracy Loss for Order-of-Magnitude Compute Savings

Combining the two: accuracy cost *A*(*m*) = *MSE*(*m*) rises slowly and monotonically with *m* (bias squared [(*m* − 1)*ρθ*]^2^grows quadratically but remains small when *m* ≪ *q*), while compute cost *C*(*m*) ∝ 1/*m* drops substantially (speedup factor *m*). This asymmetry is the value of RPS: sacrificing a small amount of accuracy yields compute savings proportional to the group size, up to order-of-magnitude (*m*-fold) levels. It is important to emphasize that there is no one-size-fits-all “optimal group size“: larger *m* saves more computation, smaller *m* yields higher accuracy, and the choice depends on the accuracy requirements or compute budget of the researcher --- compute-constrained researchers can take a larger *m* for feasibility, while accuracy-driven researchers take a smaller *m*. From the dual structure of monotonically increasing *A*(*m*) and monotonically decreasing *C*(*m*), two practical decision criteria emerge: Criterion 1 (given accuracy tolerance *τ*, most compute savings): choose the largest group size *m*^†^ = *maxm*: *MSE*(*m*) ≤ *τ* satisfying *MSE*(*m*) ≤ *τ*, maximizing compute savings without exceeding the accuracy baseline (speedup factor *m*^†^). Criterion 2 (highest accuracy given compute budget). Derive the minimum group size *m*^†^ from the affordable number of estimations, achieving the highest accuracy within the budget. These two criteria correspond to “accuracy-first” and “compute-first” scenarios, respectively. The real data analysis in Section 4 provides a concrete example: with a compute budget of approximately 24 hours (Criterion 2), we selected the data size and grouping scheme accordingly, and quantified the corresponding accuracy loss and compute savings.

## 4. Real Data Case Study

### 4.1 Data Description and Two-Layer Empirical Validation Logic

We use the GSE189050 dataset^[14]^ (a systemic lupus erythematosus (SLE) peripheral blood mononuclear cell (PBMC) CITE-seq dataset). To avoid cross-cell-type confounding, we restrict our analysis to a single cell type --- Memory B cells --- and use disease status as the binary outcome *Y* (*Y* = 1 for SLE active, *Y* = 0 for healthy controls, discarding inactive cases), yielding a subset of *n* = 2120 cells. Gene selection is performed entirely within the highly variable gene framework: we first select approximately 4096 highly variable genes using FindVariableFeatures, then randomly draw *q* = 500 as target genes (this case study is designed solely to validate the random walk statistical mechanism, not to test specific biological hypotheses, so a predefined target gene set is unnecessary; random sampling is consistent with the spirit of random grouping and avoids selection bias). The remaining highly variable genes serve as background genes. Section 2.1 has theoretically proposed the random walk characterization of grouping deviation --- this is the theoretical model to be verified (pure mathematical derivation, not empirical). This section validates it on real data using a two-layer empirical approach, making the conclusion more accessible and robust: (First empirical layer, PCA background method) using the top principal components of the background genes as DML control variables --- PCA is a closed-form, zero-iteration, near-zero-compute linear compression, used to first demonstrate that “the random walk indeed exists” with minimal computational cost; (Second empirical layer, deep learning method) using a “retrain autoencoder per group + orthogonal score” deep learning + DML pipeline to confirm the same conclusion --- deep learning achieves stronger background fitting through nonlinear representations, at the cost of expensive training computation. The two empirical methods converge to the same result, both reproducing the diffusion-dominated random walk predicted by the theoretical model, demonstrating that the conclusion is independent of the specific background compression method and thus more credible. The high training cost of deep learning vividly highlights the value of “grouping for compute savings.“

### 4.2 Random Walk Experimental Design

The two empirical layers below (PCA background method and deep learning method) share the same random walk experimental design; the two methods differ only in “how to compress the background to obtain the control variables,” so this is described first in a unified manner. Since real data has no known ground truth causal effects, we take the *m* = 1 estimate *θ^_j_*(1) (individual estimation, full control) as the gold standard for each gene (corresponding to the random walk starting point *S_j_*(1) = 0). We then progressively increase the group size, running RPS for *m* ∈ 1, 50, 100, 250, 500. Here we distinguish two different “counts“: the compute cost *C*(*m*) = *K* = *q*/*m* is the number of autoencoder training runs for a single RPS execution (*m* = 1 requires *q* runs, *m* = *q* requires only 1), serving as the horizontal axis of the accuracy-compute trade-off. To characterize the increment distribution at each step (building the diffusion/power-law picture of the random walk), we perform *R* = 20 independent random partitions for intermediate group sizes (*m* ∈ 50, 100, 250), obtaining one *S_j_*(*m*) sample per gene per repetition. *m* = 1 and *m* = *q* have unique partitions, so we run once each. Each intermediate group size thus yields *q* × *R* deviation samples, sufficient to estimate the full increment distribution. The *R* repetitions are a statistical tool, increasing total training to *R* × *C*(*m*) (corresponding to the “multiple partition averaging” recommendation in Section 5.3), without changing *C*(*m*) = *q*/*m* as the trade-off axis. The cumulative deviation *S_j_*(*m*) = *θ^_j_*(*m*) − *θ^_j_*(1); the diffusion is characterized by the variance *Var*[*S*(*m*)] across all (gene × repetition) samples (dominant feature), the drift is characterized by the mean of *S_j_*(*m*) (generally negligible), consistent with the picture that each gene is an independent random walk trajectory and the variance expands with step count. Additionally, we use three-dimensional density surfaces (horizontal axis *m*, vertical axis *S*, height = density) to visualize the broadening of the distribution with *m*.

**Table:** Group structure and autoencoder training counts for each group size (*q* = 500). The number of groups *K* = ⌈ *q*/*m*⌉ equals the number of autoencoder training runs per RPS execution (i.e., the compute cost *C*(*m*), the horizontal axis of the accuracy-compute trade-off, decreasing with *m*); the number of repetitions *R* is a statistical tool for characterizing the increment distribution (20 for intermediate *m*), with total training = *K* × *R*, summing to 841.

| Group size $m$ | Groups $K = \lceil q/m \rceil$ | Targets per group | Repetitions $R$ | AE runs per RPS = $C(m)$ | Total AE runs = $K \times R$ |
| --- | --- | --- | --- | --- | --- |
| 1 | 500 | 1 | 1 | 500 | 500 |
| 50 | 10 | 50 | 20 | 10 | 200 |
| 100 | 5 | 100 | 20 | 5 | 100 |
| 250 | 2 | 250 | 20 | 2 | 40 |
| 500 | 1 | 500 | 1 | 1 | 1 |
| **Total** | --- | --- | --- | --- | **841** |

Why *m* = 1 and *m* = 500 are run only once: Neither involves the “random partition” degree of freedom, so repetitions provide no new information. For *m* = 1, each target gene is alone in its own group with all other variables as controls --- the partition is unique, and this is the gold standard starting point of the random walk (*S_j_*(1) = *θ^_j_*(1) − *θ^_j_*(1) ≡ 0, repetition is meaningless). For *m* = 500 (= *q*), all 500 target genes fall into the same group with no out-of-group targets; each gene faces exactly the same, fixed background control set (background genes only), with no randomness in “which targets go to which group.” Although autoencoder training has random weight initialization, this is merely estimation noise under the same deterministic setting and does not constitute different random walk realizations, so one run suffices (symmetrically with *m* = 1). In contrast, for intermediate group sizes, each repetition changes which target genes are grouped together, directly altering the set of omitted variables each gene faces, thus requiring multiple samples to characterize the distribution.

### 4.3 PCA Background Method: Most Compute-Efficient Random Walk Validation

As the first empirical layer, we use the simplest, most compute-efficient linear principal component analysis (PCA) background method to validate the random walk. Given that the number of background genes far exceeds the sample size in single-cell data, we do not directly use all background genes as linear controls. Instead, we perform PCA on the “background genes + out-of-group target genes” matrix, take the top *d* = 32 principal components *Z_PCA_*^(*g*)^ as the group low-dimensional control variables, and estimate the causal effect for each target gene *T_j_* in the group using the orthogonal score equation *θ^_j_* = *mean*(*T_res_* ⋅ *Y_res_*)/*mean*(*T_res_*^2^) (linear *T* ∼ *Z_PCA_*, logistic *Y* ∼ *Z_PCA_*).

**Figure 7:**
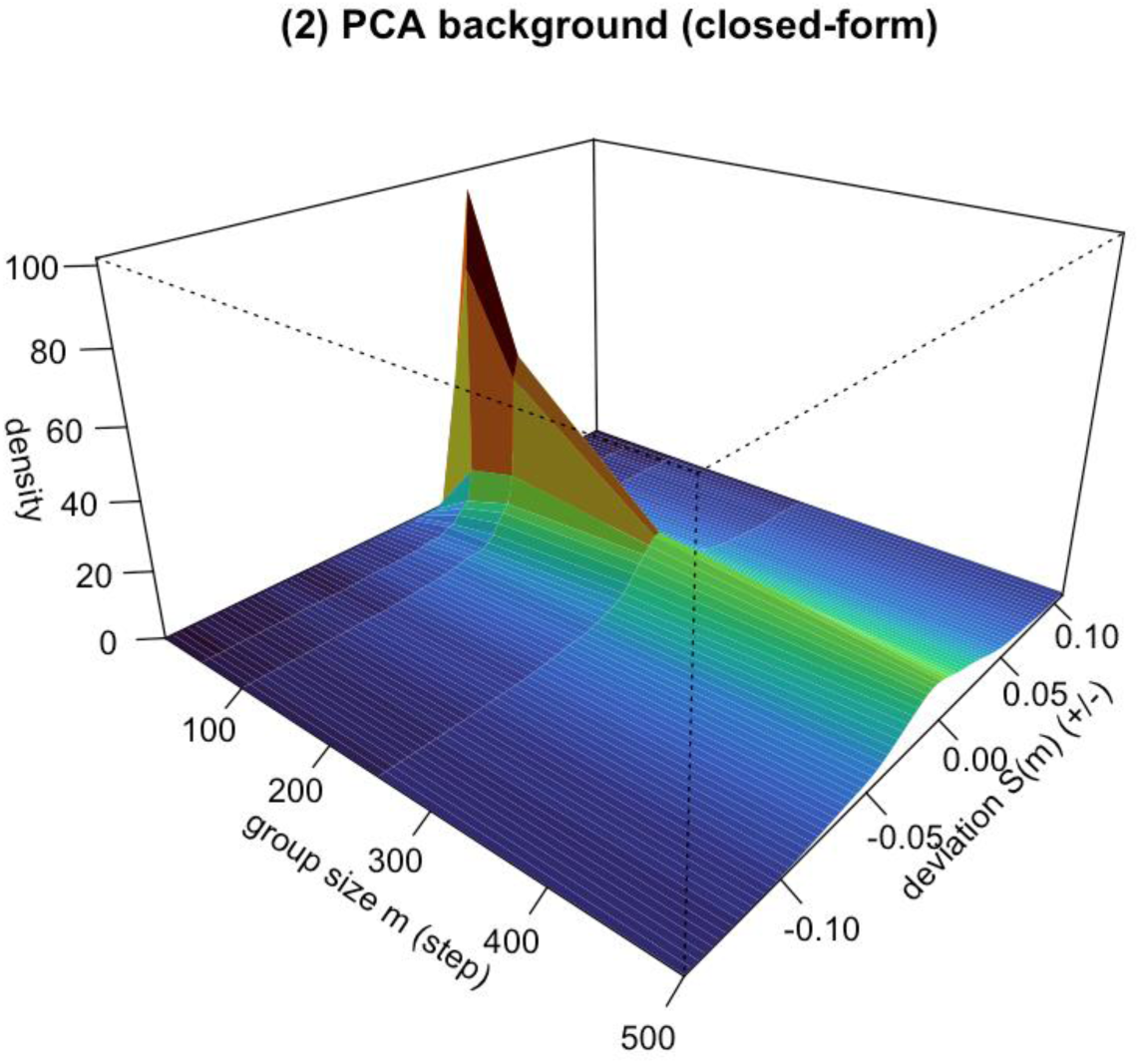
Terrain map of grouping deviation random walk for the PCA background method on real SLE data (closed-form verification of the first empirical layer, “PCA background method“; GSE189050 SLE PBMC, Memory B cells ∩ {SLE ACT, Control}, *n* = 2120 cells, *q* = 500, principal component dimension *d* = 32; intermediate group sizes *m* ∈ 50, 100, 250 with *R* = 20 random partitions each). Horizontal axis: group size *m* (the “steps” of the walk); vertical axis: cumulative deviation *S_j_*(*m*) (± both directions); height (color, sqrt scale): the probability density at that (*m*, *S*) estimated from all (gene × repetition) samples. The terrain is approximately symmetric about *S* = 0: the observed drift magnitude is only ∼ 10^−3^with positive and negative fluctuations (no systematic trend), while the diffusion *Var*[*S*(*m*)] expands monotonically with *m* (0 → 4.2 × 10^−4^ → 7.5 × 10^−4^ → 1.8 × 10^−3^ → 3.1 × 10^−3^ for *m* = 1, 50, 100, 250, 500), visually confirming the diffusion-dominated, drift-free random walk. PCA is a closed-form solution with near-zero compute, yet sufficient to reproduce this mechanism.

### 4.4 Deep Learning Method: Trading Compute for Stronger Nonlinear Representation

As the second empirical layer, we confirm the same random walk conclusion using a deep learning + DML pipeline. The deep learning background compression in this paper specifically uses an autoencoder; “deep learning method” and “autoencoder” refer to the paradigm level (contrasting with linear PCA) and the specific model, respectively, and can be used interchangeably when clear. Given the highly nonlinear nature of single-cell background expression, we depart from linear PCA and follow a two-stage pipeline. The key is that RPS divides the *q* target genes randomly into *K* = ⌈ *q*/*m*⌉ groups and retrains the autoencoder for each group: First stage (unsupervised nonlinear compression, once per group): For group *g*, input the “background genes + out-of-group target genes” (all variables except the *m* target genes in this group) into an autoencoder, take the bottleneck layer output as the *d* = 32-dimensional latent representation *Z*_ℎ*at*_^(*g*)^, with the deep network capturing the nonlinear structure of single-cell data. The *m* target genes in the group are excluded from the autoencoder input, becoming omitted confounders --- precisely the source of random walk increments; since the input changes with each group, the autoencoder must be retrained for each group. Second stage (orthogonal score estimation, DML): Using the group latent representation *Z*_ℎ*at*_^(*g*)^ as control variables, perform partialling-out for each target gene *T_j_* in the group to obtain residuals *T_res_* = *T_j_* − *E^*[*T_j_*|*Z*_ℎ*at*_^(*g*)^] (linear) and *Y_res_* = *Y* − *E^*[*Y*|*Z*_ℎ*at*_^(*g*)^] (logistic, following the binary outcome treatment in RA), then estimate the causal effect using the orthogonal score equation *θ^_j_* = *mean*(*T_res_* ⋅ *Y_res_*)/*mean*(*T_res_*^2^), which satisfies Neyman orthogonality. The compute cost is directly manifested by deep learning: autoencoder training is the most expensive step in the entire pipeline, and since it is trained once per group, its invocation count equals exactly the number of groups *K* = ⌈ *q*/*m*⌉. This makes the compute cost *C*(*m*) = *q*/*m* no longer an abstract nominal quantity, but a concrete number of deep network training runs: *m* = 1 requires *q* = 500 training runs (most expensive, confirming that individual estimation is computationally infeasible); *m* = *q* requires only 1 run (cheapest). Deep learning thus not only handles nonlinearity but also gives the compute cost a clear physical carrier as group size changes. It is worth emphasizing that the purpose of this case study is to prove that grouping deviation follows a random walk model, not to pursue an accurate causal number. Precisely because autoencoder training is expensive, the compute cost of individual estimation (*m* = 1) requiring *q* training runs is particularly prominent, and the compute savings from increasing the group size to reduce training runs to *q*/*m* are particularly considerable --- the value of “grouping for compute savings” is immediately apparent within the deep learning method itself. This highlights the core significance of shared background / RPS in trading a small accuracy loss for substantial compute savings. An unexpected finding is that although both methods converge on the random walk conclusion, the rate of accuracy deterioration differs markedly: from *m* = 50 to *m* = 500, the mean squared displacement of PCA increases by approximately 7.4 times, while that of deep learning increases by only about 16%. This may stem from how the two compression representations store information: PCA is a linear orthogonal projection; once a target gene is excluded from the control set, its weight in the linear projection is directly erased, and the information loss is discrete and cumulative. The autoencoder, in contrast, employs nonlinear distributed representations --- each bottleneck dimension simultaneously encodes the joint distribution of multiple genes --- so even when a target gene is excluded, its expression pattern may still be partially “covered” by the joint encoding of remaining genes in the bottleneck, thereby “buffering” the marginal effect of omitted confounders. In other words, nonlinear background compression is inherently more robust to omitted confounders than linear compression, consistent with the theoretical characterization of autoencoders as “nonlinear generalizations of PCA.” Another possible contributing factor is the truncation instability of PCA principal components. Although both methods use a fixed *d* = 32-dimensional representation, PCA principal components are sorted by eigenvalue magnitude and hard-truncated at the 32nd dimension. When the out-of-group target genes change with different partitions and the set of included background variables shifts accordingly, both the directions and the ordering of the principal components are re-estimated. The information near the truncation boundary (between the 32nd and 33rd components) is particularly sensitive to these changes, and the sorting jitter amplified by hard truncation injects additional *m*-dependent fluctuations into *θ^*. In contrast, the autoencoder bottleneck representation is an end-to-end optimized nonlinear mapping that does not rely on hard truncation based on variance ranking in a fixed orthogonal basis, and is more robust to input set perturbations. This also suggests that classical linear methods with predefined compression structures are more prone to introducing instability when input conditions change, while data-driven deep learning representations can partially mitigate this side effect of model assumptions. It should be noted that the above mechanistic explanations for the difference in diffusion rates remain speculative; this paper has not conducted ablation experiments (e.g., systematically varying the truncation dimension *d* or comparing different compression architectures) to rigorously distinguish between them, leaving this for future research.

**Figure 8:**
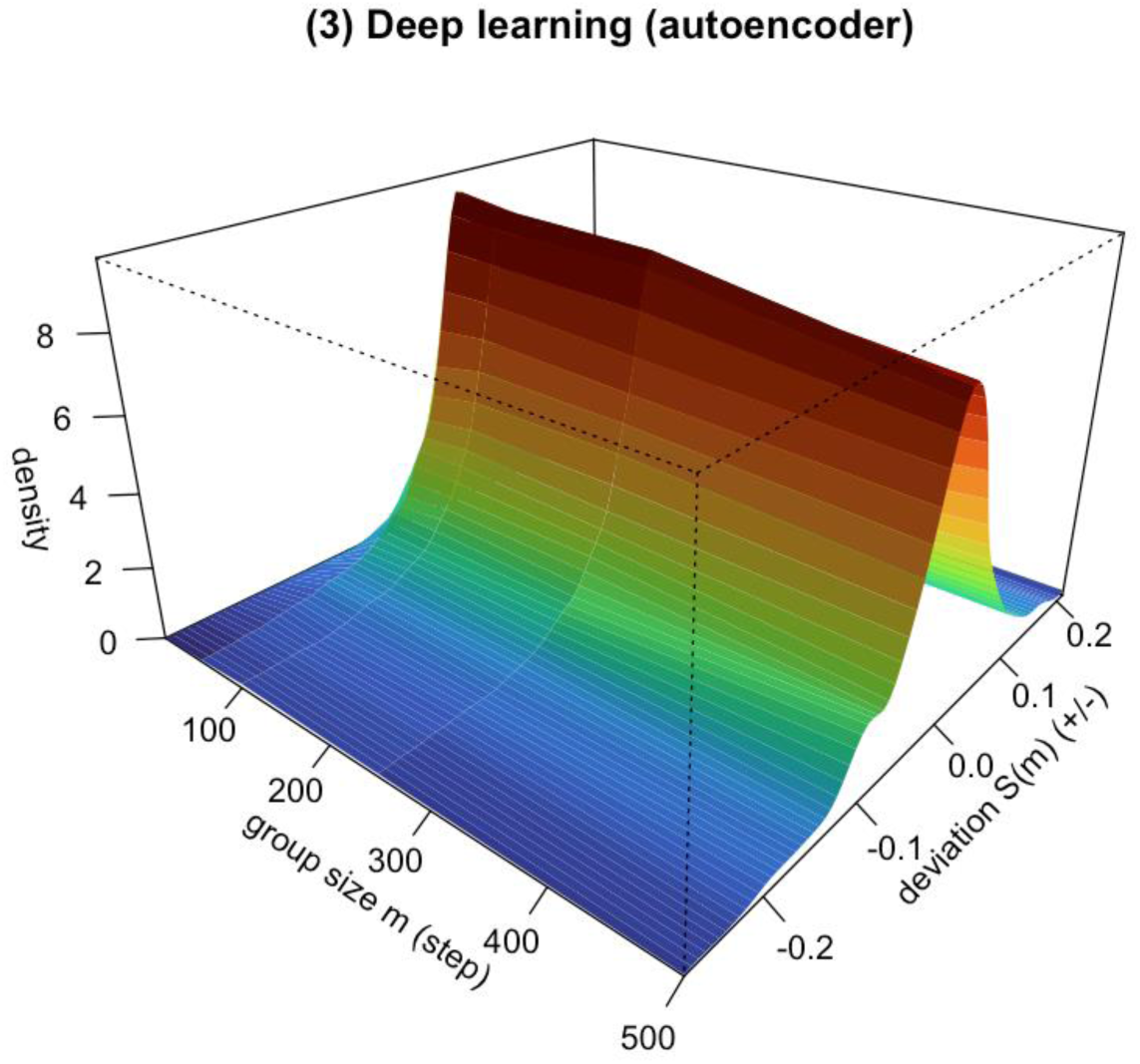
Terrain map of grouping deviation random walk for the deep learning method (autoencoder + DML) on real SLE data (nonlinear validation of the second empirical layer, “deep learning method“; GSE189050 SLE PBMC, Memory B cells ∩ {SLE ACT, Control}, *q* = 500, latent dimension *d* = 32; intermediate group sizes with *R* = 20 random partitions each). Axes are consistent with the PCA terrain map: horizontal axis group size *m*, vertical axis cumulative deviation *S_j_*(*m*), height is the probability density from all (gene × repetition) samples. Converging with the PCA background method --- similarly presenting a diffusion terrain symmetric about *S* = 0 and monotonically expanding with *m*, confirming that the random walk conclusion is independent of the specific background compression method; the only difference lies in compute: the autoencoder is retrained once per group for a total of *K* = *q*/*m* runs, while PCA is a zero-compute closed-form solution, with a striking difference in training cost between the two, highlighting the core value of grouping for compute savings.

The random walk validation results from the deep learning + DML pipeline (Fig. 9) are consistent with the theoretical predictions of Section 2.1: (1) Diffusion expands with *m* (dominant feature): the cross-(gene × repetition) variance of the deviation expands monotonically with *m*; the mean squared displacement *MSD*(*m*) = *mean S*^2^(*m*) increases monotonically with no U-shape, providing real-data evidence for “accuracy cost is monotonic, *m* = 1 is optimal“; the log-log slope of the SLE data is close to the standard diffusion value of 1, consistent with the drift-free random walk; a deviation from 1 would suggest modular co-expression (super/sub-diffusion); (2) Drift is negligible: the cross-500-gene average cumulative deviation is very small with no systematic monotonic trend --- real gene correlations and effects have mixed signs, so the increment mean *μ_j_* ≈ 0 and the drift term (*m* − 1)*μ_j_* is negligible, with the grouping deviation manifesting as a diffusion-dominated drift-free random walk; linear drift appears only in same-signed co-expression modules; (3) Compute cost: *C*(*m*) decreases monotonically as autoencoder training runs *K* = *q*/*m* (*m* = 1 requires 500 deep network training runs, *m* = 500 only 1), forming an asymmetric trade-off with the monotonically increasing *A*(*m*) = *MSD*(*m*): small accuracy loss yields order-of-magnitude compute savings. Since the most expensive autoencoder training count equals the number of groups, deep learning not only handles the nonlinear fitting of single-cell data but also gives the “grouping for compute savings” benefit a clear physical carrier: larger groups mean fewer deep networks to train, saving more computation.

**Figure 9:**
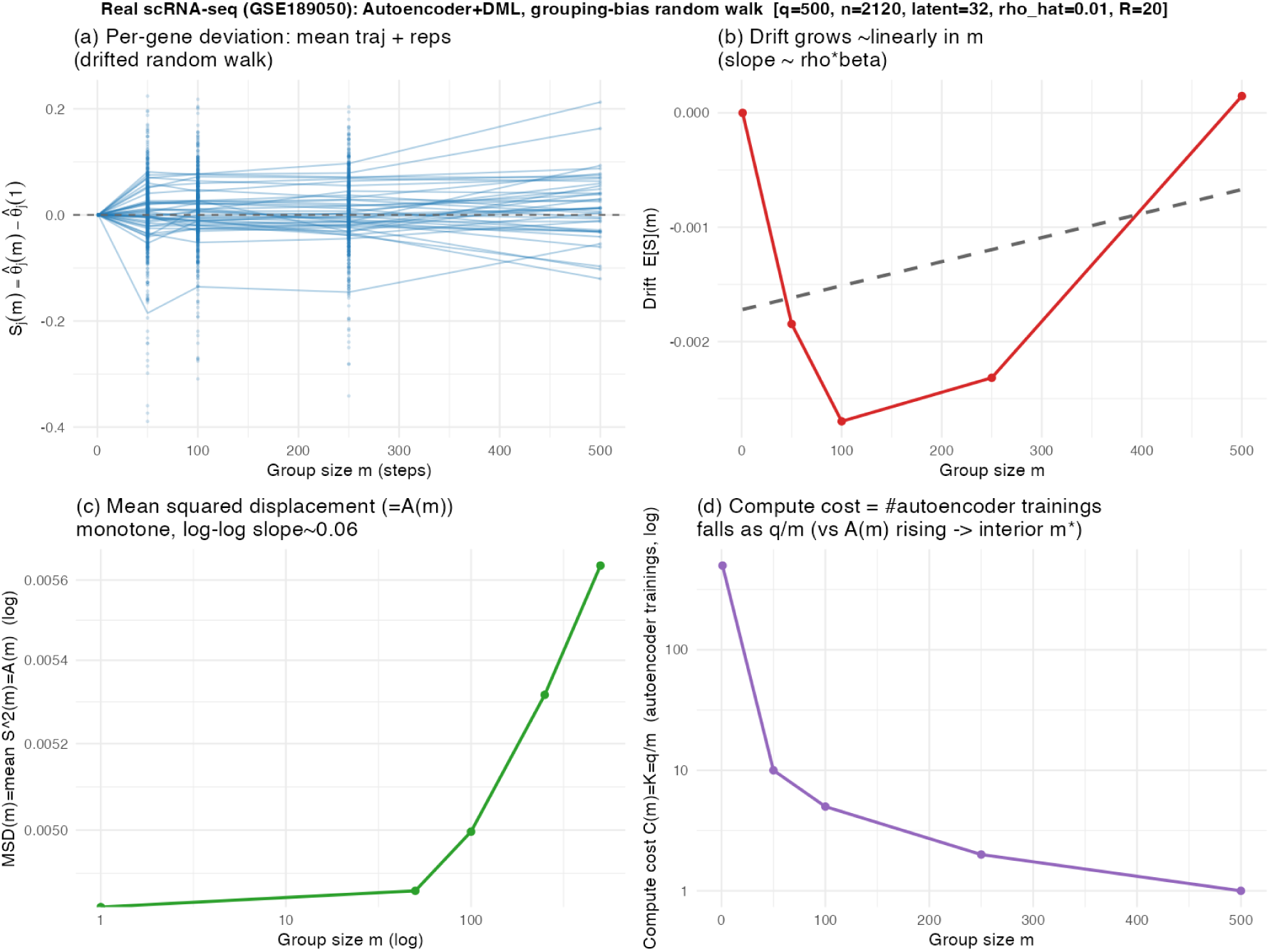
Four-panel validation of the grouping deviation random walk by “autoencoder + DML” on single-cell real data (GSE189050). (a) Cumulative deviation trajectories *S_j_*(*m*) of several genes exhibit a (generally drift-free) random walk; (b) Drift: generally negligible, small in magnitude; (c) Diffusion: mean squared displacement *MSD*(*m*) = *A*(*m*) increases monotonically, with the log-log slope characterizing the scaling exponent; (d) Compute cost *C*(*m*) = *K* = *q*/*m* decreases with *m*, forming an asymmetric trade-off with the increasing *A*(*m*).

### 4.5 Significance of the Convergent Results

The PCA background method (Section 4.3) and the deep learning method (Section 4.4) yield consistent conclusions on the same data and same experimental design: the cumulative deviation *S_j_*(*m*) from grouped estimation manifests as a diffusion-dominated random walk in both cases, with diffusion expanding monotonically with *m*, mean squared displacement increasing monotonically with no U-shaped turning point, and observed drift negligible. This method independence is itself important evidence --- the random walk is not an artifact of any specific compression method (linear or nonlinear), but rather an intrinsic statistical consequence of the grouping operation in the presence of target variable correlations, making the conclusion robust and reliable.

## 5. Discussion and Conclusion

### 5.1 Main Findings

This paper proposes the Randomized Partition Strategy (RPS), dividing *q* target genes into *K* = *q*/*m* groups, sharing one background compression per group, reducing deep learning training runs from *q* to *K* --- a factor of *m* savings. Theoretical analysis shows that the cumulative deviation *S_j_*(*m*) = *θ^_j_*(*m*) − *θ_j_* of the RPS estimator follows a drift-free symmetric random walk: diffusion 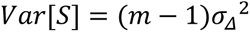 holds universally, corresponding to variance accumulation; drift *E*[*S*] = (*m* − 1)*μ_j_* appears only when the product of correlation and effect has a systematic same sign (special case). The mean squared displacement *E*[*S_j_*(*m*)^2^] = *MSE*(*m*), so the accuracy cost is predictable. Under the DML setting, *MSE*(*m*) is a monotonically increasing convex function of *m* with no interior minimum --- *m* = 1 (individual estimation) is always optimal. Accuracy cost rises slowly while compute cost drops substantially (*C*(*m*) ∝ 1/*m*, speedup factor *m*), forming an asymmetric trade-off. On the GSE189050 SLE single-cell data, the two methods PCA and DL converge to the same conclusion, validating that the random walk mechanism is independent of the specific compression method. Plain-language summary: In simple terms, the core finding of this paper can be understood as follows --- if you imagine the causal estimate for each target gene as a “guessing game,” then each time another gene shares the same background information with it (while that gene itself is not “controlled”), it is equivalent to injecting a random perturbation into that guess. The more genes that share, the more perturbations accumulate, but the perturbation direction is random (positive and negative cancel out), so the overall guess does not systematically bias in any direction --- it simply “wobbles” over a range that expands with the number of shared genes. This is the random walk. More importantly, the expansion of this wobble range is predictable (proportional to the number of steps), so researchers can know in advance: how much accuracy will be lost for a given group size, and how much compute will be saved.

### 5.2 Limitations and Future Directions

Grouping sensitivity under high-correlation modules: The closed-form theoretical derivation in this paper adopts the homogeneity assumption (equal pairwise correlations and equal effects among all target variables), but real data typically exhibits modular correlation structures --- genes within the same pathway or co-expression module are highly correlated, while correlations between modules are low. Based on random grouping, our randomly drawn target genes have a very low average correlation (observed *ρ^* ≈ 0.01), so the bias increment from omitted group-mates is small and the random walk is gradual. However, in clinical and single-cell practice, researchers often focus on highly correlated genes within the same pathway or co-expression module; if such genes are placed in the same group and simultaneously excluded from background control, then the omitted confounders are precisely these strong confounders, and the bias increment *ρ_jk_ θ_k_* accumulates rapidly, potentially causing severe accuracy loss. This suggests that the good performance of random grouping implicitly relies on the premise of “low within-group correlation“: when the target set itself is highly correlated, accuracy loss should be carefully evaluated, or correlation-aware grouping should be employed --- distributing strongly correlated genes across different groups (so that within-group omitted confounders are as uncorrelated as possible, keeping strongly correlated variables as mutual background controls), rather than placing them in the same group. How to design optimal grouping when the correlation structure is known is a valuable theoretical question.

### 5.3 Practical Recommendations

The Randomized Partition Strategy is most valuable when compute becomes the practical bottleneck, particularly when the goal is “global screening for signals” --- first screen with grouped estimation, then refine the few hits with *m* = 1. The practical accessibility of this method: even on a personal computer (with GPU), using deep learning as the background compression method together with the RPS strategy, it is feasible to complete initial screening of DML causal inference effect estimates for hundreds or even thousands of genes within a reasonable time.

### 5.4 Conclusion

This paper proposes the Randomized Partition Strategy (RPS), dividing *q* target genes into *K* = *q*/*m* groups, sharing one background compression per group, reducing deep learning training runs from *q* to *K* --- a factor of *m* savings. Theoretical analysis and real-data validation (GSE189050 SLE single-cell data, *n* = 2120) demonstrate that the cumulative deviation *S_j_*(*m*) = *θ^_j_*(*m*) − *θ_j_* follows a drift-free symmetric random walk, with diffusion variance ∝ (*m* − 1) expanding monotonically and mean squared displacement equal to the mean squared error, so accuracy loss is predictable. Two practical conclusions follow: (1) *m* = 1 is always optimal, accuracy cost is monotonically increasing convex with no U-shaped turning point; (2) a small accuracy loss yields *m*-fold compute savings, with group size flexibly chosen based on accuracy requirements or compute budget --- first screen with grouped estimation, then refine the few hits with *m* = 1. This work provides a quantifiable theoretical foundation for compute strategy selection in single-cell high-dimensional causal inference.

## Ethical Approval statement

This study used publicly available datasets. In accordance with relevant institutional and regulatory guidelines, publicly available datasets do not require ethical review and approval. Therefore, no ethical approval from an institutional review board was necessary for this study.

